# Complex genetics cause and constrain fungal persistence in different parts of the mammalian body

**DOI:** 10.1101/2022.01.20.477007

**Authors:** Martin N. Mullis, Caleb Ghione, Michael Lough-Stevens, Ilan Goldstein, Takeshi Matsui, Sasha F. Levy, Matthew D. Dean, Ian M. Ehrenreich

## Abstract

Determining how genetic polymorphisms enable certain fungi to persist in mammalian hosts can improve understanding of opportunistic fungal pathogenesis, a source of substantial human morbidity and mortality. We examined the genetic basis of fungal persistence in mice using a cross between a clinical isolate and the lab reference strain of the budding yeast *Saccharomyces cerevisiae*. Employing chromosomally-encoded barcodes, we tracked the relative abundances of 822 genotyped, haploid segregants in multiple organs over time and performed linkage mapping of their persistence in hosts. Detected loci showed a mix of general and antagonistically pleiotropic effects across organs. General loci showed similar effects across all organs, while antagonistically pleiotropic loci showed contrasting effects in the brain and the kidneys, liver, and spleen. Persistence in an organ required both generally beneficial alleles and organ-appropriate pleiotropic alleles. This genetic architecture resulted in many segregants persisting in the brain or in non-brain organs, but few segregants persisting in all organs. These results show complex combinations of genetic polymorphisms collectively cause and constrain fungal persistence in different parts of the mammalian body.

## Introduction

Fungi are a major class of opportunistic human pathogen, infecting billions and killing millions of people per year ^1,2^. Hundreds of diverse fungal species are known to infect humans ^3^. These fungi mainly infect the immunocompromised, an increasing segment of the global population due to improvements in medicine that have lowered the mortality associated with life-threatening conditions ^3^. Such opportunistic infections can be difficult to treat ^2–5^, but the identification of mechanisms enabling fungi to cause these infections may facilitate the development of more effective antifungal therapies ^6–12^.

Many opportunistic fungal pathogens can be challenging to work with genetically ^13,14^ However, the budding yeast *Saccharomyces cerevisiae,* one of the main eukaryotic model organisms in biology, is also an opportunistic pathogen, with numerous isolates obtained from clinical infections ^10–12,15–17^. Humans are regularly exposed to *S. cerevisiae*, as it occurs naturally in the environment and is used in the production of beer, wine, bread, chocolate, and other foods and dietary supplements ^18–22^. Notably, *S. cerevisiae* is in the same Saccharomycetaceae family of Ascomycete yeasts as *Candida*, the main genus involved in opportunistic fungal infections (*Candida albicans*)^23^.

The ability to infect immunocompromised humans varies among *S. cerevisiae* strains ^11^, with clinical isolates found throughout the species’ genealogy ^17,24^ Despite their lack of genetic relatedness, clinical *S. cerevisiae* isolates are thought to possess similar traits, including the ability to attach to and penetrate surfaces and tolerance to human body temperature ^10,11,17,21,25,26^. However, determining why certain strains are able to infect humans ultimately requires mapping the specific genetic polymorphisms that cause opportunistic pathogenicity and determining the traits they affect. Such work is challenging because it requires performing genetic mapping in yeast inside mammalian hosts, such as mice.

The most powerful genetic mapping approach in *S. cerevisiae* is linkage mapping with large mapping panels of haploid meiotic progeny (segregants) ^27,28^. When two haploid isolates are crossed and sporulated, their haploid segregants each receive a unique random combination of alleles from their parents ^29^. This shuffling of genetic material makes it possible to measure traits of interest in segregants and then link these traits to specific genomic locations (loci) that cause phenotypic differences ^30^. Examination of large numbers of segregants provides the statistical power to identify loci explaining most of the heritable differences in traits of interest ^27,28^.

To enable linkage mapping with an *S. cerevisiae* cross in mice, we generated a panel of haploid segregants with known genotypes and chromosomally-encoded barcodes ^28,31^, which were genetically engineered for high-throughput phenotyping as a pool. We mated the lab reference strain (BY) and a haploid derivative of the 322134S clinical isolate (3S)^24^. BY and 3S are highly diverged at the sequence level, with a genetic difference present every ~270 base pairs ^32,33^. While 3S is an isolate obtained from the throat sputum of a patient with a clinical infection ^24^, BY is a commonly used reference strain that is avirulent and rapidly cleared by mice ^10^.

We used this BYx3S cross to obtain insights into how genetic differences among strains influence the persistence of yeast in mice. This was done by injecting a pool of 822 barcoded BYx3S *MATα* segregants into the mouse bloodstream and enumerating segregant abundances in host organs over time by barcode sequencing. Linkage mapping with these data identified numerous loci that explained most of the heritable variation in yeast persistence in hosts. Some loci showed consistent effects across host organs (general loci), while others had counteracting effects across host organs (antagonistically pleiotropic ^34–36^ loci), causing different segregants to be superior in different organs. Our work advances the use of *S. cerevisiae* as a model for opportunistic fungal pathogenicity and host-microbe interactions.

## Results

We crossed haploid BY and 3S strains that were genetically engineered to produce segregants amenable to pooled, high-throughput phenotyping. *FLO11* and *FLO8*, which respectively encode the main cell surface flocculin in this organism ^37,37,38^ and its primary transcriptional activator ^39^, were deleted from these strains. These deletions eliminate cell clumping and flocculation within and between segregants, which are problematic for pooled experiments. However, they also diminish surface adhesion and invasion, limiting our insight into these traits. BY and 3S were also engineered to have a genomic landing pad at the neutral *YBR209W* locus, enabling site-specific integration of barcodes into segregants ^28,31,40^.

The engineered BY and 3S strains were mated to produce a BY/3S diploid, which was sporulated. To ensure balanced allele frequencies and random multilocus genotypes among segregants, we performed tetrad dissection and randomly chose and barcoded one *MATα* haploid from each of 822 tetrads using transformation with a random barcode library ^28,31,40^ (**Supplementary Fig. 1**). 86 segregants were marked with two additional distinct random barcodes and these replicates were included in our experiments. Illumina sequencing was used to genotype segregants and determine their barcodes. All barcoded segregants and replicates were then grown individually to stationary phase and combined into a single pool in equimolar fractions.

Thirty-six mice were infected with 1×10^7^ cells from the segregant pool by tail vein injection (**Fig. 1a**). Equal numbers of male, female, immunocompromised (injected with 4 mg/mL dexamethasone [dex]), and immunocompetent (injected with water) mice were included. No morbidity or mortality was observed. One-third of the mice were euthanized at each of three time points (one, two, and five days post-injection). From each mouse, we harvested the brain, gonads, kidneys, liver, and spleen. The five organs were dissected, homogenized, and plated on selective media to isolate yeast from the mouse cells. On average, 69,150 colony forming units (CFU) were recovered per liver, 32,032 CFU per spleen, 3,843 CFU per kidney pair, 1,741 CFU per brain, and 69 CFU per gonad pair. For every organ, recovery decreased over time, suggesting clearance of at least some segregants by the mice (**Fig. 1b**). Recovery was lowest in the brain and gonads, which both have blood barriers ^48,49,57^.

**Figure 1.**
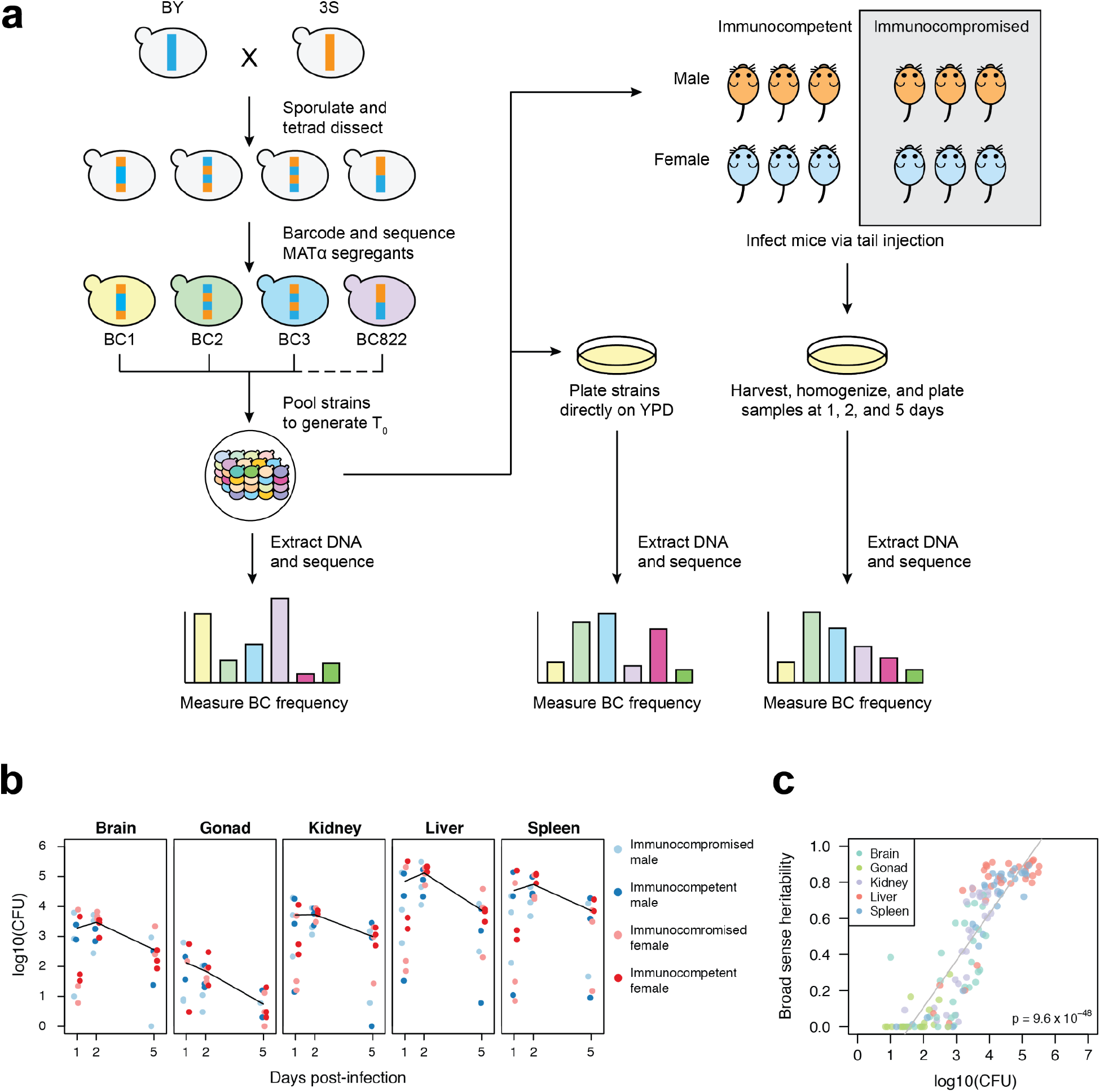
Experimental infections of mice using a pool of barcoded, haploid segregants. (**a**) The workflow and general design of the experiment. Haploid BY and 3S strains were crossed, the resulting diploid was sporulated, and tetrads were dissected to generate a panel of 822 recombinant haploid progeny. These strains were then barcoded with unique 20mer nucleotide sequences and pooled together (T_0_). T_0_ was used to infect mice using tail vein injection and simultaneously plated in triplicate onto control plates containing rich medium. At three time points post-infection, organ samples were collected from infected mice and plated on rich medium. After two days of growth on plates, DNA was extracted from the yeast and sequenced to measure relative barcode abundance of each strain. DNA was also extracted from T_0_ to measure initial barcode frequencies. For each strain, the change in normalized barcode frequency relative to T0 after correcting for on-plate growth was used as the phenotype in all downstream analysis. (**b**) Plot reporting the CFU recovered from each organ sample across time points. Each panel shows the samples from a different organ. Color of each point corresponds to the sex and immunological state of the mouse from which the sample was recovered. Mean log_10_(CFU) over time is shown as a black line. (**c**) Scatterplot showing the broad-sense heritability (H^2^) of each organ sample as a function of the CFU recovered from that sample. The color of each sample corresponds to organ type.

Barcode sequencing was performed on all samples. For every barcode in each sample, we divided the change in barcode frequency between the time of sampling and the initial pool by the initial barcode frequency. A linear model was then used to correct these changes in barcode frequency for differences in growth among segregants in on-plate controls (**Supplementary Fig. 2**). Residuals from this correction for on-plate growth were used as persistence phenotypes in downstream analyses. Of 180 processed samples (five organs x two sexes x two dex treatments x three timepoints x three replicates), yeast were present in 166 and recovered from 139; only these samples were analyzed further.

We used the 86 segregants that were replicated in the pool to estimate broad sense heritability (H^2^) in each sample. Across samples, H^2^ ranged from 0 to 0.92 (median H^2^ = 0.57). Our ability to measure H^2^ was strongly affected by yeast recovery from samples, with higher recovery resulting in higher H^2^ (simple linear regression of H^2^ on CFU, R^2^ = 0.77, p = 9.6×10^−48^; **Fig. 1c**). Variability in yeast recovery among samples was presumably due to both differences in clearance among mice and organs, as well as technical factors associated with organ dissociation. Despite limitations associated with recovering yeast from dissociated organs, the high H^2^ values in many samples shows that genetic polymorphisms among segregants caused differences in persistence in hosts.

To distinguish samples with significant differences in persistence among segregants, we applied one-way analysis of variance (ANOVA) to each sample, again using the replicated segregants. 94 samples showed significant differences among segregants (Bonferroni-corrected α = 0.05 threshold, p ≤ 3.7×10^−4^). Of these, two were excluded because they had distorted measurements for persistent segregants, suggesting their sequencing libraries were of low quality (**Supplementary Fig. 3**). Only a single gonad sample showed significant differences in persistence among segregants; this sample, which had a lower H^2^ value (0.29), was also omitted from later analyses due to a lack of organ replicates (**Supplementary Fig. 4**). In the 91 remaining significant samples from the brain, kidneys, liver, and spleen, the median H^2^ was 0.75 (from 0.24 to 0.92), indicating most of the variability among segregants in these replicated, higher quality samples was genetic in origin (**Fig. 1c**).

We analyzed the relationships among the 91 samples using hierarchical clustering, principal components analysis (PCA), and examination of pairwise correlations. All methods found the same result: the samples split into two clusters, brain and non-brain (kidneys, liver, and spleen) (**Figs. 2a** and **b**; **Supplementary Fig. 5**). In PCA, these groups were visible in the loadings on the first principal component (PC_1_), which was the only PC to account for a meaningful portion of the variance across samples (54.1%; other PCs explained ≤7.4% of the variance across samples; **Supplementary Table 1**). Whether a sample was from the brain or a non-brain organ was the only experimental factor showing a major relationship with PC_1_, explaining 85.2% of the variance in PC_1_ in a one-way ANOVA (p = 8.84×10^−39^). Time and the interaction between brain vs. non-brain and time also had minor significant effects, each explaining <2.7% of the variance in PC_1_ (full-factorial ANOVA with brain vs. non-brain, time, and brain vs. non-brain-time interaction, factor effect test p < 2.94×10^−3^). Such time effects would be expected if selection acts on phenotypic differences among segregants and these differences vary across organs. Immunological state and sex showed no relationship with PC_1_, perhaps because we directly injected yeast into the bloodstream. Following these results, we generated aggregate brain and non-brain measurements for each segregant by averaging data from the 15 brain samples and 76 non-brain-samples, respectively. These aggregate measurements showed a poor but highly significant correlation (Spearman’s rho = 0.21, p = 1.78×10^−9^; **Fig. 2c**).

**Figure 2.**
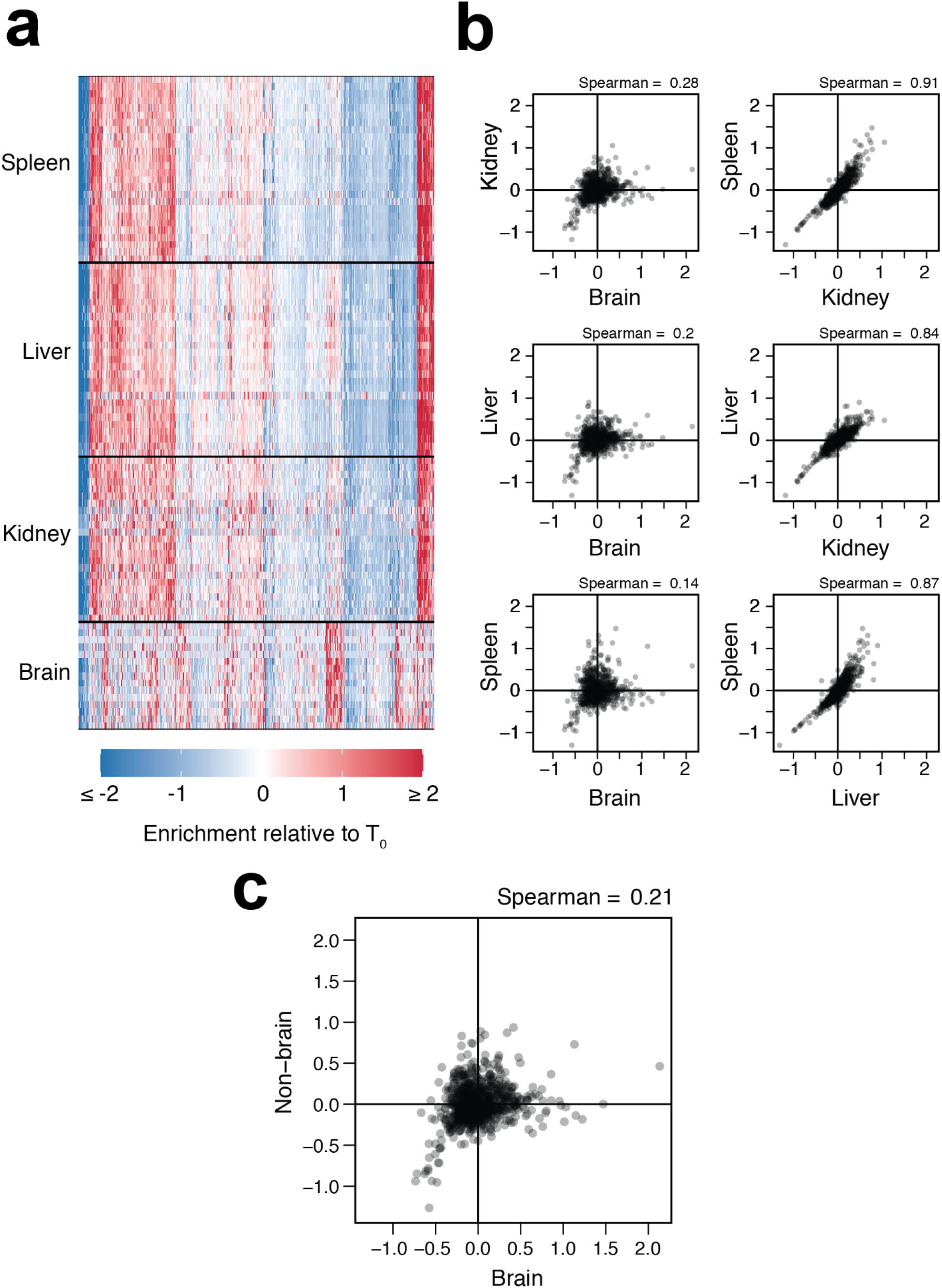
Organ type is the main driver of variation in persistence across samples. (**a**) Heatmap showing phenotypes of strains (x-axis) across organ samples (y-axis). Samples are clustered by organ type and segregants are clustered by phenotype across non-brain samples. (**b**) All pairwise comparisons of segregant phenotypes between organs. Here, each segregant phenotype in an organ represents it’s average measurement across all samples for that organ in which reproducible loci were detected. (**c**) Comparison of segregants’ aggregate phenotypes in the brain and non-brain samples.

These above results show that segregants show reproducible differences in persistence in brain and non-brain organs, and indicate that the genetic bases of persistence in these different parts of the host body are likely only partially overlapping. We began determining the genetic basis of these differences in persistence within and between organs. We performed linkage mapping on each of the samples with significant phenotypic differences among segregants, detecting 494 loci in total (**Fig. 3a**; **Supplementary Table 2**). On average, 5.43 loci were identified per sample (min: 0, max: 12) and 90 of 91 samples had at least one detected locus that was also identified in another sample. Multiple loci were mapped in the spleen (181), liver (157), kidney (101), and brain (55), and many loci were detected in numerous samples (min = 2, max = 87, median = 4), as expected if segregants show reproducible phenotypes across samples due to a common set of loci. These detections could be consolidated to 35 distinct loci, based on overlapping confidence intervals (**Supplementary Table 3**). The number of loci identified in these samples showed a highly significant relationship with H^2^ (simple linear regression of number of loci on H^2^ R^2^ = 0.32, p = 4.02×10^−9^; **Supplementary Fig. 6**), suggesting heterogeneity in measurement noise across samples impacted statistical power.

**Figure 3.**
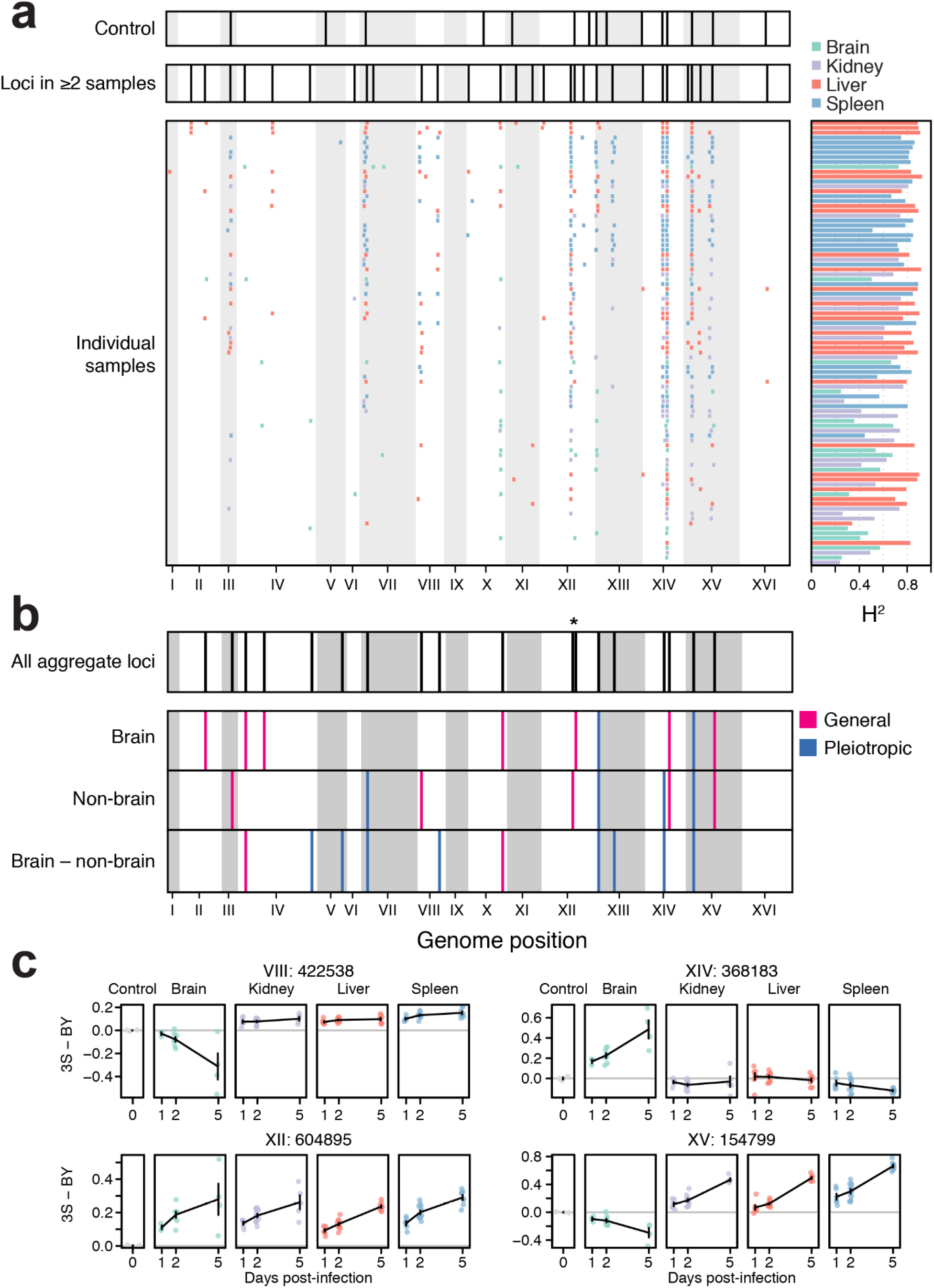
Identification of loci associated with persistence in the host in organ samples. (**a**) Consolidated loci in the plate controls (*top*), consolidated loci across all organ samples (*middle*), and individual loci detected in each organ sample (*bottom*) shown in descending order from greatest to least number of loci detected. Corresponding broad sense heritability (H^2^) measurements for each sample are shown to the right of each individual sample. Samples are colored by organ type. (**b**) Loci detected in genome-wide scans using aggregate data across samples (*top*), followed by loci detected using mean segregant phenotypes in brain and non-brain samples, as well as the difference in mean phenotype between brain and non-brain samples (*bottom*). Red loci have effects of the same sign in both brain and non-brain samples (general), while blue loci have effects with opposite signs in brain and non-brain samples (pleiotropic). * indicates two linked, but distinguishable, loci on chromosome XII. (**c**) The effects of loci detected in whole-genome scans using aggregate data from the brain samples and non-brain samples, as well as the difference between brain and non-brain samples, are shown. Effects were calculated as the mean persistence of strains with the 3S allele at the focal locus minus the mean persistence of strains with the BY allele after correction foron-plate growth. The effect of each locus in the controls after correcting for on-plate growth is also shown (gray).

To improve statistical power and minimize the chance of false positives, we performed linkage mapping on aggregate brain and non-brain measurements, as well as the difference between the two; these measurements should be more precise than data from individual samples. The scans on brain, non-brain, and difference measurements respectively identified nine, nine, and 10 loci (**Fig. 3b**). Some loci were mapped in multiple of these scans, resulting in the identification of 18 unique loci. Seven of these loci overlapped loci detected in controls, but in these cases loci showed different effects between the samples and controls (**Figs. 3a** through **c**; **Supplementary Fig. 7**). Loci detected in the brain, non-brain, and brain vs. non-brain scans explained 89.7%, 62.3%, and 83.6% of H^2^ in their respective measurements.

The resolution of loci was poor, with confidence intervals from scans using aggregate data spanning 58 kb (min: 9 kb, max: 100 kb) and 32.5 genes (min: 6, max: 62) on average **Supplementary Table 4**). To better resolve these loci, we leveraged confidence intervals from detections of these loci in multiple individual samples. While average resolution was only slightly improved (mean interval = 24 kb, mean number of genes = 12, min number of genes = 2, max number of genes = 39; **Supplementary Table 5**). The two most finely resolved loci were each localized to two candidate protein-coding genes ^41^. A locus on Chromosome XIV spanned a subunit of the BLOC-1 complex involved in endosomal maturation (*SNN1*) and a poorly understood, pleiotropic gene (*MKT1*) known to influence many quantitative traits in *S. cerevisiae* ^42–45^. A locus on Chromosome XV encompassed alcohol dehydrogenase (*ADH1*) and a gene regulated by phosphate levels (*PHM7*). Additionally, a locus on Chromosome XII fractionated into two distinct intervals, one including only the genes for DNA topoisomerase III (*TOP3*) and a thiamine transporter (*THI7*).

We next focused on understanding how the 18 loci influence the ability of segregants to persist in different parts of the mammalian body. We calculated the effects of each locus in the brain and non-brain organs. Loci were then classified as general or antagonistically pleiotropic if the same allele or different alleles were superior in both the brain and non-brain organs, respectively (**Fig. 4**). Of the 18 loci, ten were general and eight showed antagonistic pleiotropy between the brain and non-brain organs. Eight of the beneficial alleles at the general loci were contributed by the 3S clinical isolate (**Fig. 4a** upper right quadrant), as opposed to two by the BY lab strain (**Fig. 4a** lower left quadrant). By contrast, eight loci showed antagonistic pleiotropy between the brain and non-brain organs, suggesting a fitness trade-off between persistence in the brain and other organs (**Figs. 4a**, **c**, and **d**).

**Figure 4.**
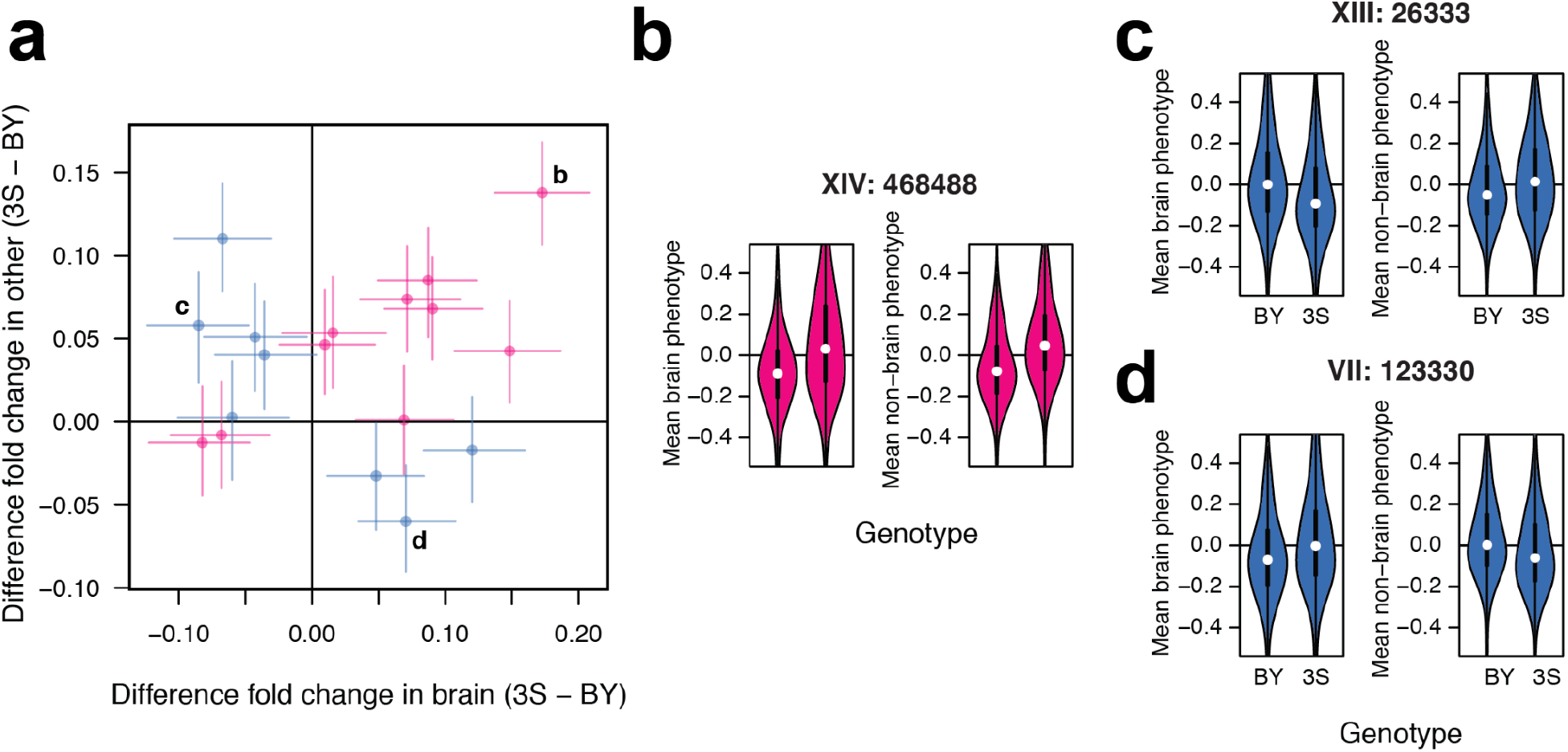
Identified loci show a mixture of general effects and antagonistic pleiotropy. (**a**) The effect sizes of loci detected using aggregate phenotype data in the brain and non-brain organs are shown on the x- and y-axes, respectively. Positive effect sizes mean that strains carrying the 3S allele were enriched in the samples while negative values mean that strains carrying the BY allele were enriched. Loci are colored by whether the same allele is beneficial in both brain and non-brain samples (red; general effects) or not (blue; antagonistic pleiotropy). Numbers correspond to specific examples in **b** and **c**. (**b**) A locus with a general effect on persistence within the host. Brain (left) and non-brain (right) phenotypes are plotted as a function of strain genotype at this locus. Positional information for the locus is denoted by bold text above the example. (**c**) An antagonistically pleiotropic locus at which the BY allele is beneficial in the brain (left) and detrimental in other organs (right). (**d**) An antagonistically pleiotropic locus at which the 3S allele is beneficial in the brain (left) and detrimental in other organs (right).

We determined how alleles at general and antagonistically pleiotropic loci combine to cause fungal persistence. We found a positive relationship between the number of beneficial alleles at general loci and persistence in both brain and non-brain organs (simple linear regression of number of general alleles on persistence, brain R^2^ = 0.19 and p = 9.79×10^-39^, non-brain R^2^ = 0.1 and p = 7.29×10^-21^; **Fig. 5a**). Similarly, we found that the number of brain or non-brain alleles at antagonistically pleiotropic loci was positively and negatively related to segregants’ persistence in the brain (simple linear regression of number of brain alleles on brain persistence, R^2^ = 0.12, p = 1.32×10^-23^) and non-brain organs (simple linear regression of brain alleles on non-brain persistence, R^2^ = 0.08, p = 2.51×10^-16^), respectively (**Fig. 5b**).

**Figure 5.**
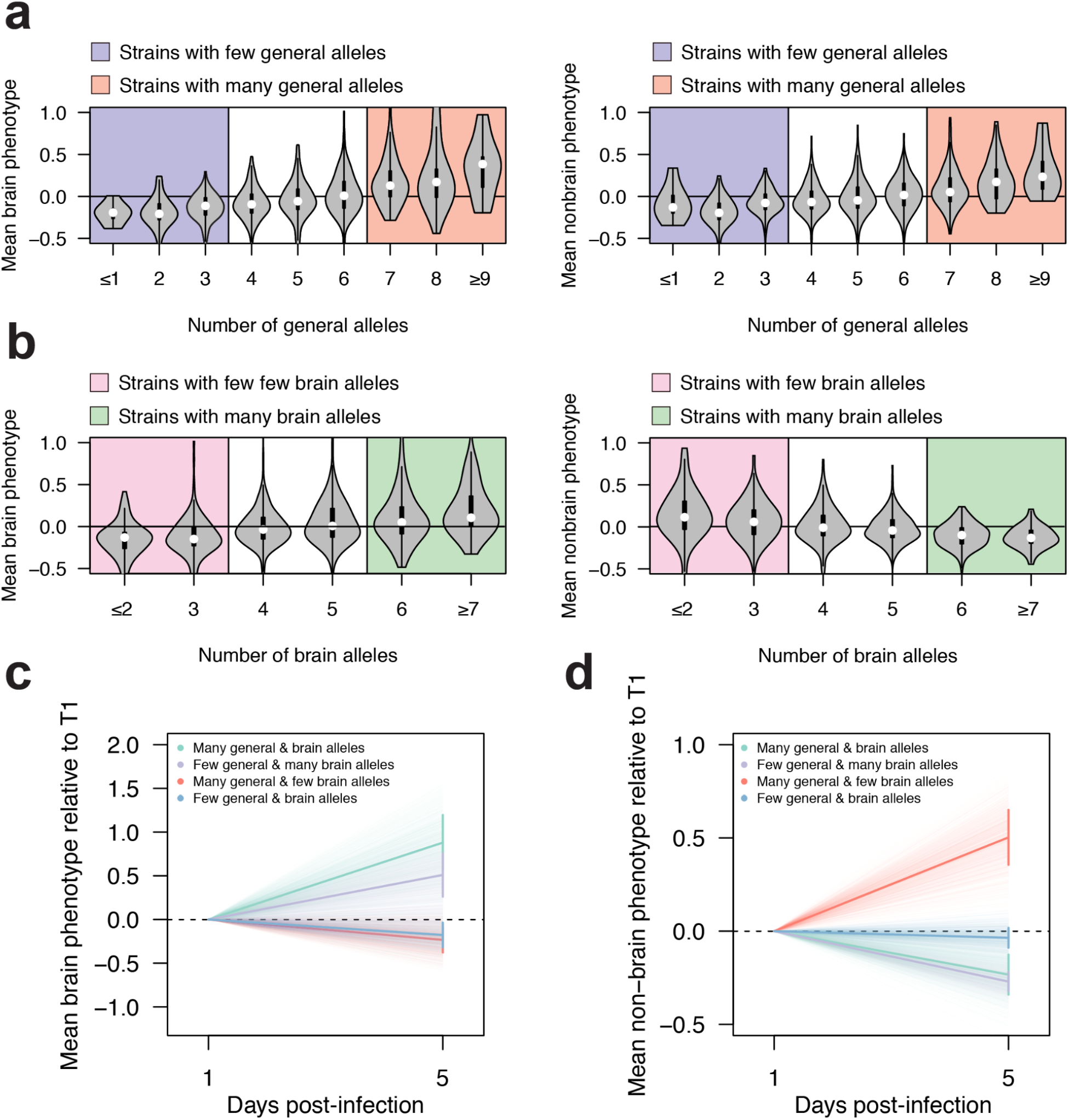
General and antagonistically pleiotropic loci collectively influence strain persistence in the host. (**a**) Violin plots showing the mean strain phenotypes in the brain samples (left) and non-brain samples (right) as a function of the number of generally beneficial alleles present in a segregant. Thresholds for strains considered to have a high or low number of general persistence alleles are represented by colored backgrounds. (**b**) Violin plots showing the mean strain phenotypes in the brain (left) and non-brain samples (right) as a function of the number of antagonistically pleiotropic brain alleles present in a strain. Thresholds for strains considered to have a higher or low number of alleles favoring persistence in the brain over other organs are represented by colored backgrounds. (**c**) Plot showing mean change in enrichment in the brain samples over time relative to T_1_ measurements (bold lines) for strains that have a high or low number of generally beneficial alleles as well as a high or low number of alleles favoring persistence in the brain over other organs (according to thresholding in panels **a** and **b**). Error bars show the standard error about the mean enrichment of strains at five days post-infection. Faint lines show the enrichment over time of bootstrapped data (1,000 replicates). (**d**) Plot showing mean enrichment in the non-brain samples over time relative to day one measurements (bold lines) for strains that have a high or low number of generally beneficial alleles as well as a high or low number of alleles favoring persistence in the brain over other organs (according to thresholding in panels **a** and **b**). Error bars show the standard error about the mean enrichment of strains at five days post-infection. Faint lines show the enrichment over time of bootstrapped data (1,000 replicates).

Lastly, we examined how sets of general and antagonistically pleiotropic loci jointly influence persistence. In a given organ, the most persistent segregants were enriched for both the beneficial alleles at general loci and the appropriate alleles at antagonistically pleiotropic loci (brain 2×2 *χ*^2^ test: *χ*^2^ = 12.39, p = 4.3×10^-4^; non-brain 2×2 *χ*^2^ test: *χ*^2^ = 17.36, p = 3.1×10^-5^; Figs. 5c and d; Supplementary Table 6). In the brain, segregants were even able to persist if they were enriched for organ-specific alleles alone, although their persistence was lower than segregants enriched for both organ-specific and generally beneficial alleles (Fig. 5c).

## Discussion

We used barcode sequencing to phenotype a pool of genotyped segregants in mice. Analysis of segregants replicated in the pool showed that persistence in mice has a largely genetic basis, and comparison of samples revealed different segregants are superior in the brain and in the kidneys, liver, and spleen. Although technical noise limited our ability to map many loci in individual samples, aggregating brain and non-brain samples made it possible to identify loci explaining most of the variability in persistence within and between organs. 18 loci were detected, with the majority having effects across all organs. Some of these loci were generally beneficial, while others exhibited antagonistic pleiotropy, showing tradeoffs between brain and non-brain organs. These antagonistically pleiotropic loci could represent either single polymorphisms that have different effects in distinct parts of the host body or closely linked polymorphisms with different effects.

Our findings may explain why diverse *S. cerevisiae* isolates act as opportunistic pathogens ^17,24^. The ability to persist in mammalian hosts is highly polygenic: we identified 18 loci in a cross of two isolates and examination of additional isolates would likely detect even more ^46,47^. With so many loci involved, many *S. cerevisiae* isolates will possess beneficial alleles at some general loci, as we saw with both BY, an avirulent isolate ^10^, and 3S, a clinical isolate ^24^ Furthermore, all isolates will carry alleles of antagonistically pleiotropic loci that are beneficial somewhere in the host body. Thus, the mixing of genetic material throughout the species by chance outcrossing events may produce strains that can persist in particular mammalian organs. Supporting such a possibility, clinical isolates are often highly heterozygous diploids that likely resulted from recent mating events in nature ^17,26^.

Our results, in particular the identification of numerous antagonistically pleiotropic loci, also indicate different organs in the mammalian body represent distinct environments for fungi. The brain and non-brain organs have a myriad of functional and physiological differences: for example, the brain has its own semi-permeable barrier ^48,49^ and the kidneys, liver, and spleen filter blood ^50^. Persisting in these distinct organs may require different traits, which may be beneficial in some organs and detrimental in others. If these traits vary across strains, which seems likely based on our data, many individuals may only be able to infect certain organs in the mammalian body and may be constrained in their potential to spread to other organs post-infection.

Genetic mapping has the potential to help reveal molecular mechanisms shaping the abilities and constraints of persistence in different parts of the host body. Although our resolution was coarse in most cases, a few finely resolved loci implicated a potential diversity of cellular processes, including endosome maturation (*SNN1*), ethanol production (*ADH1*), genome stability (*TOP3*), phosphate metabolism (*PHM7*), and thiamine uptake (*THI7*). The other gene in these intervals (*MKT1*) has an unclear function. Notably, *ADH1* ^51^, *MKT1* ^52^, and *PHM7* ^53^ have been found to affect pathogenicity in other fungi, and both endosomal function ^54^ and thiamine transport ^55^ have been linked to virulence as well. Our system provides an opportunity not only to identify new mechanisms underlying persistence in hosts, but also to study how both known and unknown mechanisms act in combination.

Finally, fungal infections in the brain and central nervous system (meningitis) are a leading cause of morbidity and mortality among immunocompromised patients ^3,56^. We detected specific allele combinations that allowed segregants to persist in the brain, but our limited mapping resolution precluded insight into how these alleles act mechanistically. A possibility is they influenced passage through the blood-brain barrier, as we recovered fewer yeast from the brain than the kidneys, liver, or spleen. This hypothesis, which requires future testing, illustrates how our experimental system can be used to understand the mechanisms by which genetic polymorphisms modify interactions between yeast cells and the mammalian body.

## Acknowledgements

We thank members of the Ehrenreich, Dean, and Levy labs, as well as James Boedicker, Steven Finkel, and Sergey Nuzhdin, for feedback on drafts of this manuscript. We also thank Joseph Hale and Rachel Schell for helping to construct the panel of BYx3S segregants.

## Animal subjects

All procedures and personnel involving mice were approved under the University of Southern California’s Institute for Animal Care and Use Committee protocol #21102.

## Author contributions

M.N.M., C.G., M.L.S., S.F.L., M.D., and I.M.E. conceptualized this project. M.N.M. and T.M. constructed and genotyped the yeast strains. M.N.M., C.G., M.L.S., and I.G. performed the experiments. M.N.M. analyzed the data. M.N.M. and I.M.E. wrote the manuscript, incorporating input from all other authors.

